# Self-organization controls expression more than abundance of molecular components of transcription and translation in confined cell-free gene expression

**DOI:** 10.1101/401794

**Authors:** P.M. Caveney, R. Dabbs, G. Chauhan, S.E. Norred, C.P. Collier, S.M. Abel, M.L. Simpson

## Abstract

Cell-free gene expression using purified components or cell extracts has become an important platform for synthetic biology that is finding a growing numBer of practical applications. Unfortunately, at cell-relevant reactor volumes, cell-free expression suffers from excessive variability (noise) such that protein concentrations may vary by more than an order of magnitude across a population of identically constructed reaction chambers. Consensus opinion holds that variability in expression is due to the stochastic distribution of expression resources (DNA, RNAP, ribosomes, etc.) across the population of reaction chambers. In contrast, here we find that chamber-to-chamber variation in the expression efficiency generates the large variability in protein production. Through analysis and modeling, we show that chambers self-organize into expression centers that control expression efficiency. Chambers that organize into many centers, each having relatively few expression resources, exhibit high expression efficiency. Conversely, chambers that organize into just a few centers where each center has an abundance of resources, exhibit low expression efficiency. A particularly surprising finding is that diluting expression resources reduces the chamber-to-chamber variation in protein production. Chambers with dilute pools of expression resources exhibit higher expression efficiency and lower expression noise than those with more concentrated expression resources. In addition to demonstrating the means to tune expression noise, these results demonstrate that in cell-free systems, self-organization may exert even more influence over expression than the abundance of the molecular components of transcription and translation. These observations in cell-free platform may elucidate how self-organized, membrane-less structures emerge and function in cells.

## Introduction

Cell-free gene expression using purified components or cell extracts has increasingly become a viable platform for synthetic biology. Cell-free synthetic biology applications include prototyping gene circuit elements^1,2^, viral biosensors^3^, enzymes^4^, and characterizing gene regulatory elements of non-model microbial hosts^5^. Beyond these immediate applications, a broader goal is the realization of ever more complex cell-free systems^6^ that may ultimately approach cell-like capabilities^7^. However, these higher aspirations are stymied by the inability to achieve reproducible behavior from seemingly identical cell-free expression reactors, especially at cell-relevant reactor volumes^8–11^. Indeed, even in simple, single-gene, expression experiments, protein concentrations may vary by more than an order of magnitude across a population of identically constructed reaction chambers^12,13^. The consensus opinion from previous work is that stochastic distribution of expression resources (RNAP, ribosomes, etc.) across the population of reaction chambers leads to the broad distribution in the produced protein concentration^12,14,15^ (stochastic seeding hypothesis; Fig. 1A). This view would hold that identical chambers with exactly equivalent concentration of resources would produce a very narrow distribution of protein (Fig. 1B). However, this hypothesis has not been rigorously tested, perhaps due to the experimental difficulty of removing the stochasticity from the distribution of expression resources to cell-sized reaction chambers^16^.

**Figure 1.**
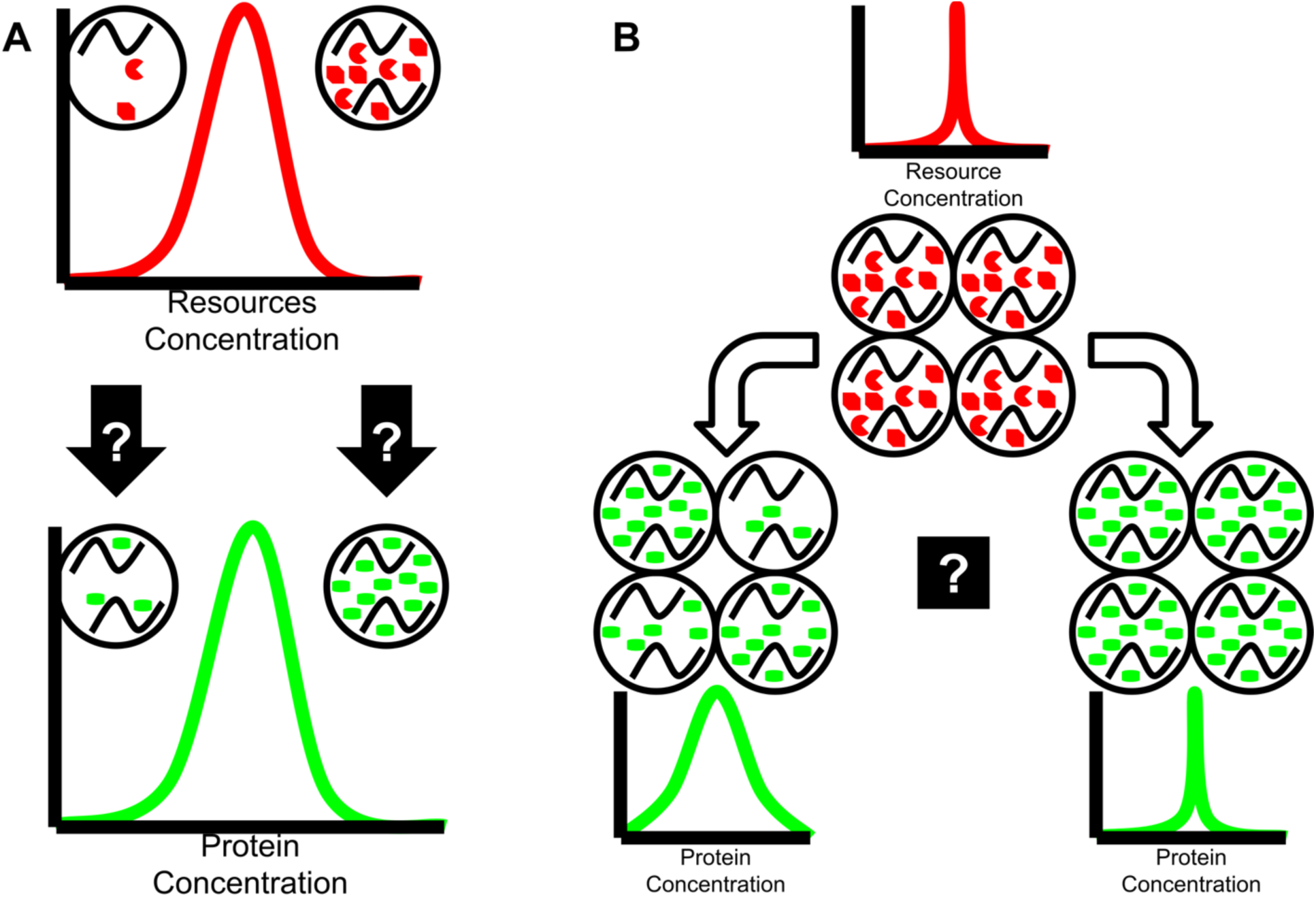
(A) (Upper) Expression resource (represented in red here) concentration varies in vesicles due to the stochastic capture of molecules during vesicle formation. (Lower) The stochastic seeding hypothesis holds that variability in protein production in vesicles is the direct result of resource concentration variation. (B) If the resource concentration hypothesis were true, a population of vesicles identical in size and resource concentration would lead to low variability in protein production (right). Conversely, high variability in protein production (left) is not consistent with the resource concentration hypothesis.

The stochastic seeding of expression resources is most often modeled as binomial selection of plasmid DNA and protein synthesis components from a well-mixed, bulk reaction mixture^16,17^. Plasmid DNA typically has the fewest number of encapsulated copies, followed by proteins (e.g. polymerase, ribosome), and finally small molecules (e.g. nucleotides, amino acids) with the greatest abundance. Accordingly, the expected encapsulated distribution is most broad for DNA and most narrow for small molecules, so it is often assumed that the DNA distribution is responsible for the variation in the expressed protein concentration^14,15,18^. Support for this hypothesis is found in the correlation between the amount of DNA and the amount of protein made in bulk (i.e. large volume) cell-free reactions^2,19^. Yet surprisingly, over a broad range of DNA concentrations, there is no^12^, or at best little^13,20^, correlation between the amount of encapsulated DNA and the amount of protein produced in reactions confined to cell-like volumes (lipid vesicles, water in oil droplets). This puzzling result and a recent review of measured and expected extrinsic noise in confined, cell-free expression^16^ cast doubt on the stochastic seeding hypothesis while suggesting that spatial effects (e.g. reactor volume) may play an important role in expression variability. Cell-free expression experiments in cell-sized containers^8–10^ and superresolution imaging of cells^21,22^ are elucidating the importance of spatial effects in gene expression behavior^23,24^. The cell studies show that membrane-less spatial organization plays important roles in gene regulation in prokaryotes^25^ and perhaps eukaryotes^26^, and recent work demonstrates that the same may be true for cell-free expression in confined and crowded environments^8,9,27^.

Here we show that self-organization of expression centers controls expression more than the abundance of the molecular components of transcription and translation in confined, cell-free synthetic biology. In cell-free expression confined in vesicles, we found that variability in protein expression was largely unrelated to the stochastic seeding of expression resources. Instead, this expression noise emerged from vesicle-to-vesicle variation in how expression resources within each vesicle were distributed across sets of expression centers (i.e. regions of correlated gene expression). Vesicles that organized into many expression centers, each having relatively few expression resources, exhibited high expression efficiency and high protein concentration. Conversely, vesicles that organized into just a few centers where each center had an abundance of resources, exhibited low expression efficiency and low protein concentration. A particularly surprising finding was that dilution of expression resources reduced the vesicle-to-vesicle variation in protein production. Vesicles with dilute pools of expression resources were more likely to exhibit high expression efficiency and low expression noise than those with more concentrated expression resources. The results presented here are similar to recent experimental results that show suB-regions in *E. coli* control the location and rates of gene expression^21^. Likewise, regulated phase transitions may induce similar suB-regions in eukaryotic cells^28^. The cell-free platforms reported here provide a flexible experimental system for understanding the emergence of self-organized structure in gene expression in confined volumes, and may offer insights into how these features arise and function in cells.

## Results and Discussion

To explore the relationship between initial expression resource concentration and final expressed protein concentration, we tracked cell-free expression of Yellow Fluorescent Protein (YFP) from pEToppYB plasmids and the PURE expression system confined in POPC vesicles (Fig. 2A; Methods). In addition, each vesicle contained a population of fluorescent molecules (AF647 conjugated to transferrin) that were captured at the time of vesicle formation. Fluorescent images were obtained using a confocal microscope to track protein (YFP) expression and AF647 fluorescence for three hours (Fig. 2B; Methods). The diameter of each vesicle was recorded to allow calculation of the reaction volume. Time traces were truncated to 90 minutes to allow noise analysis of expression^29–31^. As seen in previous experiments^32,33^, the YFP fluorescent intensity increased over time before reaching a plateau that was indicative of protein synthesis stopping, not equilibrium between protein synthesis and decay^30,34^. The AF647 signal decreased exponentially with time as the fluorophore was photobleached (Fig. 2B inset). Photobleaching over the course of the experiment reduced AF647 fluorescence 37.3% and YFP fluorescence 24.8%. The experiments produced a distribution of encapsulated AF647 at the beginning of the experiment that was indicative of the expected distribution of expression resource concentrations across the population of vesicles (Fig. 2C). The distribution of the final YFP fluorescence intensities was indicative of the distribution of the final expressed protein concentrations across the population of vesicles (Fig. 2C).

**Figure 2.**
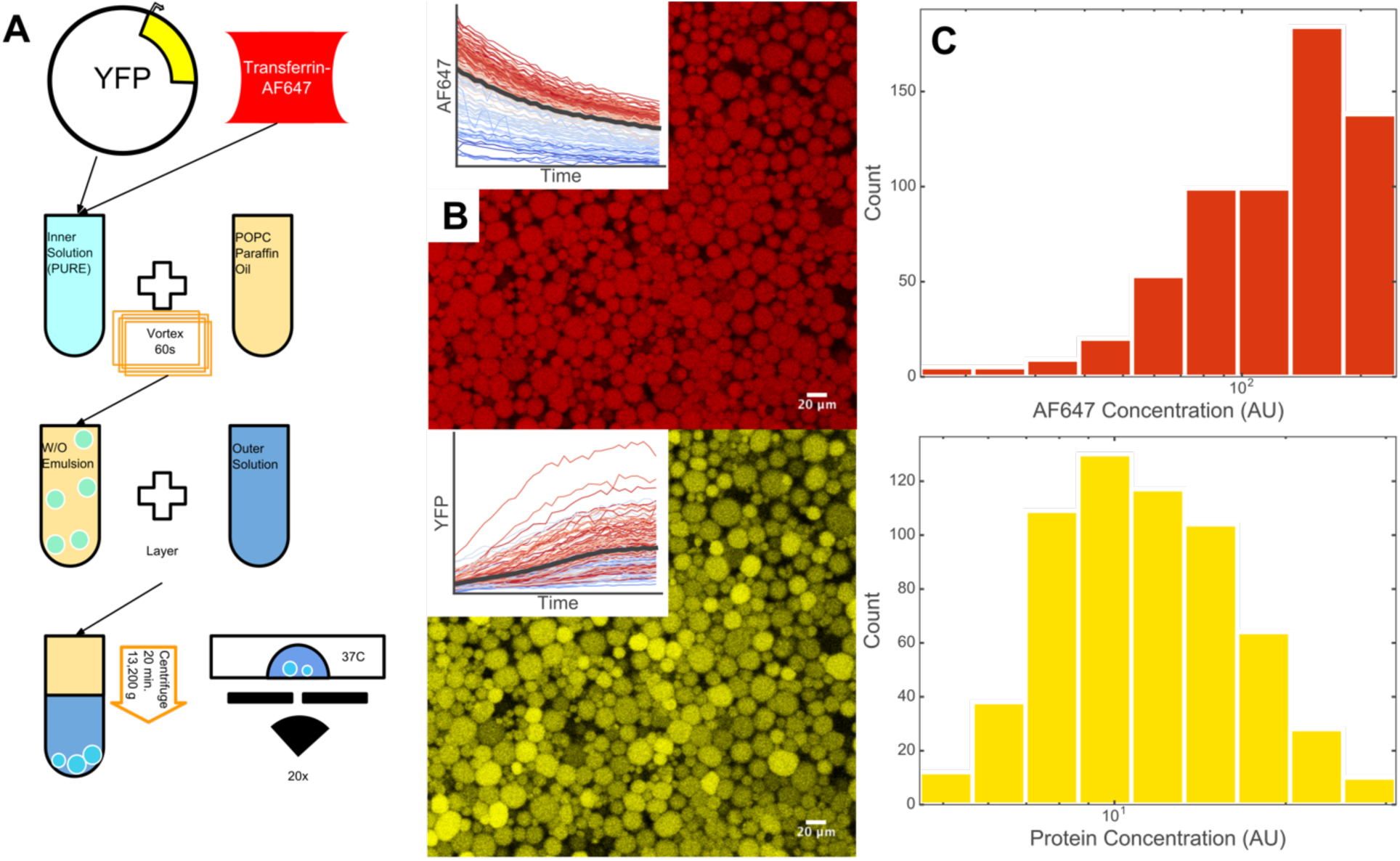
(A) YFP expressing vesicles were formed using water-in-oil emulsion technique. The PURE system, pEToppYP (YFP expressing plasmid), and AF647 conjugated to transferring were encapsulated in POPC vesicles. Vesicles were imaged at 37° C. (B) AF647 (top) and YFP (Bottom) fluorescence were tracked with a 20x air objective for 3 hours. AF647 signal photobleached (inset) while YFP signal increased (inset) before reaching a plateau. (C) Distributions of average AF647 fluorescent intensity per vesicle at the beginning of the experiment and the distribution of average YFP fluorescent intensity at the end of the experiment.

We performed experiments using 3 different concentrations of the PURE expression system that covered a 3-to-1 concentration range (1x, 0.5x, 0.33x). The 0.5x and 0.33x concentrations were made by adding nuclease-free water to the 1x PURE solution. For each PURE system concentration, we measured distributions of final YFP concentration (Fig. 3A). The mean final YFP concentration went down linearly as expression resources were diluted (Fig. 3C). While all three PURE system concentrations resulted in relatively broad YFP concentration distributions, the coefficient of variation (CV=standard deviation/mean) of these distributions decreased as the expression resources were diluted (CV of 0.38, 0.28, and 0.20 for 1x, 0.5x, and 0.33x, respectively). Strikingly, there was significant overlap between the YFP distributions resulting from the different PURE dilutions (Bhattacharyya distance between 1x and 0.5x = 0.83; between 1x and 0.33x = 0.57; and between 0.5x and 0.33x = 0.88). Many of the resource poor (0.33x) vesicles resulted in higher YFP concentrations than many of the resource rich (1x) vesicles (Fig. 3A). To ensure these effects were not related to the relatively broad distribution of vesicle sizes (∼100-3300 µm^3^), we also considered a subset of the vesicles with a narrow range of vesicle volumes (500-700 µm^3^) and observed essentially the same behavior as seen with the entire vesicle population (SI Fig. 1).

**Figure 3.**
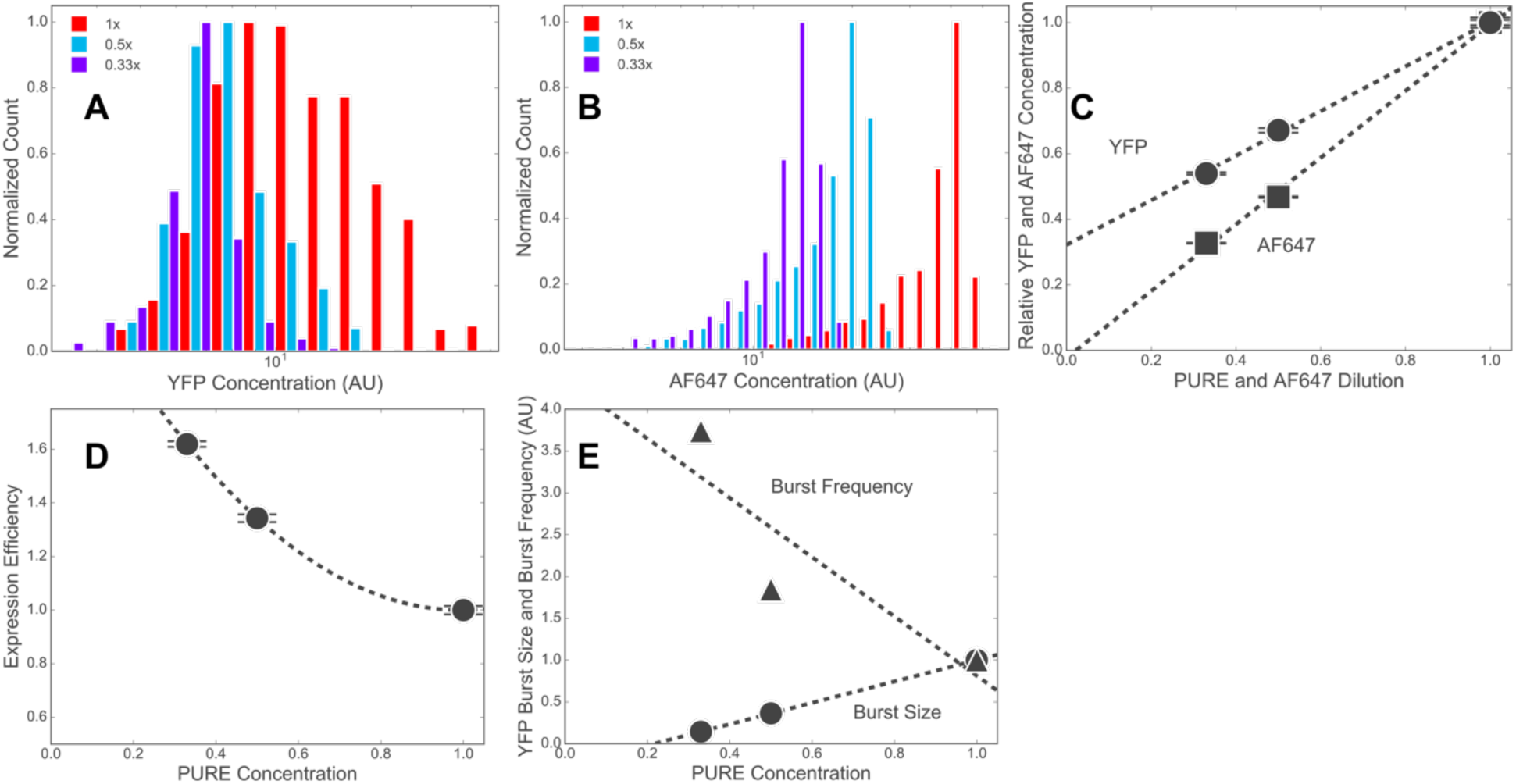
(A) Normalized distributions of final YFP concentrations in populations of vesicles with 1x (Red), 0.5x (Blue) and 0.33x (Purple) expression resource concentrations. (B) Normalized distributions of AF647 concentrations in populations of vesicles with 1x (Red), 0.5x (Blue) and 0.33x (Purple) AF647 concentrations. (C) Population average of fluorescent intensity for YFP and AF647 as the PURE system or the AF647 respectively were diluted compared to the 1x condition. Error bars are standard error of the mean. (D) Expression efficiency as the PURE system was diluted. Error bars are standard error of the mean. (E) Expression burst size and burst frequency as the PURE system was diluted. Lines are linear fits.

To quantify how the distribution of expression resources concentrations varied with the dilution of the PURE system, we performed experiments using 3 different concentrations (also 1x, 0.5x, and 0.33x) of the AF647 fluorescent marker (Fig. 3B). Like the mean final YFP concentrations, the mean concentration of AF647 went down linearly with dilution (Fig. 3C). In contrast to the YFP distributions, the AF647 distributions did not narrow as the AF647 was diluted (CV=0.25, 0.28, and 0.24, respectively for 1x, 0.5x, and 0.33x). Furthermore, compared to the YFP distributions, there was much less overlap in the AF647 distributions (Bhattacharyya distance between 1x and 0.5x = 0.37; between 1x and 0.33x = 0.18; between 0.5x and 0.33x = 0.66). In addition, to confirm previous reports^12,13,20^ that variations in the amount of plasmid DNA captured in vesicles had little effect on the amount of protein produced, we varied the concentration of the pEToppYB plasmid from 0.1x to 10x and saw no correlation between DNA and final YFP concentrations (SI Fig. 2).

These dilution experiments demonstrated that YFP expression (mean and noise) behavior was quite different than the resource concentration behavior. The dilution of resources by a factor of 3 (0.33x compared to 1x) only reduced the mean protein population by a factor of 1.85 (Fig. 3C). Furthermore, as indicated by the AF647 measurements, dilution did not change the width of the resource concentration distributions (Fig. 3B), yet significantly narrowed the YFP distribution (Fig. 3A). With this lack of correlation between resource and expressed protein concentrations, the stochastic seeding of expression resources cannot be the driving force behind the large variation in the protein concentrations. Instead, the results here suggest that there is a large vesicle-to-vesicle variation in how efficiently resources are used. Many of the resource poor (0.33x) vesicles were efficient enough to express more protein than inefficient, resource rich (1x) vesicles (Fig. 3A). Because of this large vesicle-to-vesicle variation in expression efficiency, even a hypothetical population of identical vesicles with exactly the same concentration of expression resources would lead to a broad distribution of protein production (Fig. 1B).

Interestingly, expression efficiency (E; defined here as 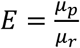,where *µ*_*r*_ = mean resource concentration and µ_*p*_= final YFP concentration) increased as resource concentration decreased (Fig. 3D). Furthermore, as expression became more efficient, the YFP distributions narrowed (i.e. noise was reduced). The reduced noise in YFP – even as the YFP concentration dropped – is at odds with the typical behavior where 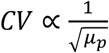. The implication is that dilution of expression resources changed expression in a way that inverted the relationship between the mean and the variance of YFP concentration, which is most often modeled as^8,31,35,36^

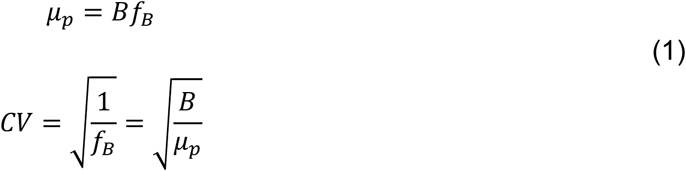

where B and *f*_*B*_ are parameters that describe the expression pattern. In the 2-state model of expression bursting from an individual gene – the episodic process where protein is produced in bursts separated by periods of little or no expression – B is the burst size (average number of protein molecules created per burst) and f_*B*_ is the burst frequency (number of bursts per unit time^29,37^). It is possible for CV to decrease even as the protein concentration drops if there is a large enough drop in the B parameter.

Using the equations above, burst size (B = *µ*_*p*_*CV*^3^; also known as the Fano factor) and 56^7^ frequency 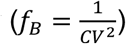 calculated from the YFP concentration distributions. Intriguingly, burst size increased but frequency decreased with increasing resource concentration (Fig. 3E). To confirm this burst size and burst frequency relationship, we also examined the noise in the 90-minute time trajectories of expression in individual vesicles using our previously reported methods^8,9^ (Methods). The burst size and frequency obtained by temporal analysis gave similar results to those found from the final YFP concentration distributions (SI Fig. 3).

In contrast to bursting from a single gene, in the multi-plasmid system described here, the burst frequency may be thought of as the number of statistically independent expression centers (Fig. 4A). An expression center might be defined by a single plasmid, or conversely, may arise from correlated expression from a group of plasmids and their associated mRNA molecules. The burst size may be thought of as the average intensity of expression from each of these centers – i.e. the average number of protein molecules synthesized from an individual mRNA or correlated ensemble of mRNA. Intuitively, one could think of an individual vesicle self-organizing into a number of expression centers (Fig. 4A) where expression in the centers are uncorrelated with each other. The number of expression centers within a vesicle determines the burst frequency, while the activity level of the bursting centers sets the burst size (Fig. 4A). High burst frequency and low burst size indicates a vesicle that sparsely distributed expression resources over a large number of expression centers. Conversely, low burst frequency and high burst size indicates a vesicle that concentrated resources in just a few centers. In studies related to the results reported here, we recently reported how spatial effects such as confinement^8^ and macromolecular crowding^9^ affect burst size and burst frequency. These reports support a model where the statistically independent expression centers are spatially distinct regions where many resources are primarily associated with a gene and/or spatially localized mRNA population. It is possible that expression centers are structurally similar to transcription factor hubs^22^ in Drosophila embryos or super-enhancers proposed in eukaryotes^26^ and functionally similar to separate resource pools in *E.coli*^25^.

**Figure 4.**
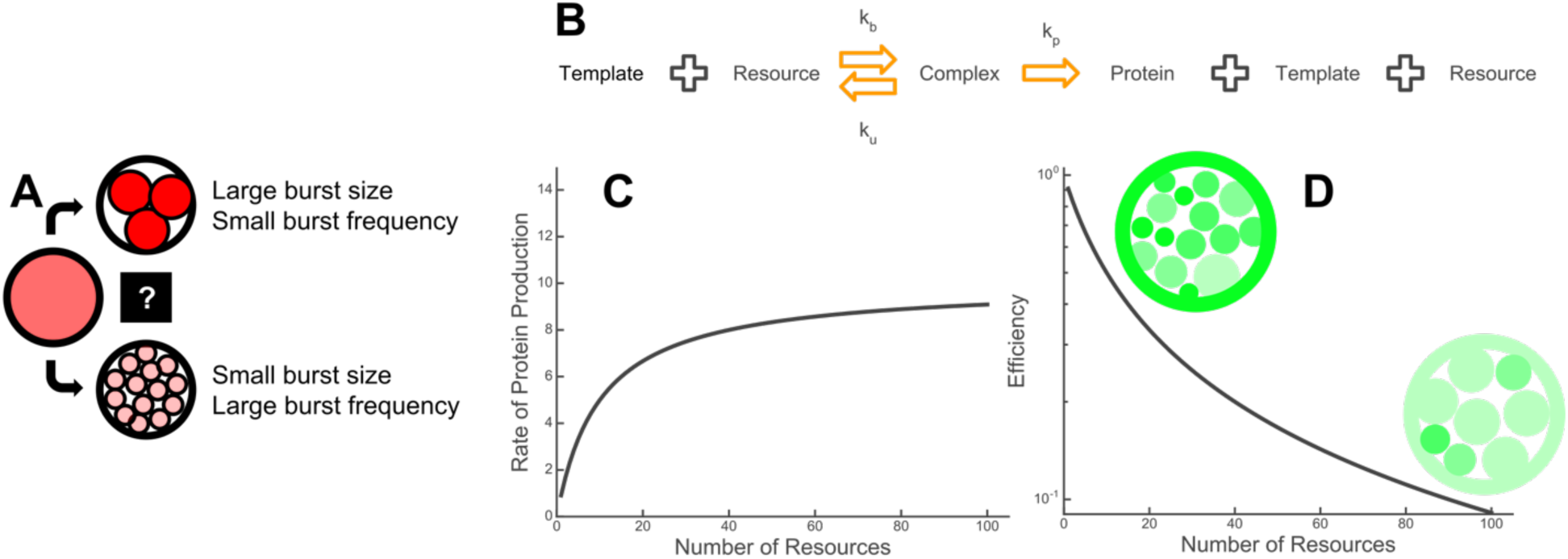
(A) A vesicle self-organizes into a number of expression domains. At the extremes, a vesicle may organize into a small number of expression domains each rich in resources, or a large number of domains each poor in resources. (B) A model of an individual domain with a single template reversibly binding resources to produce protein. (C) The rate of protein production (Michaelis–Menten model) asymptotically approaches the maximum rate as the number of resources increases. (D) The efficiency of protein production decreases as the number of resources increases.

Importantly, the results presented here demonstrate that the expression pattern selected by an individual vesicle determines its expression efficiency. To explore how the distribution of resources affected expression efficiency, we built a model of expression from an individual expression center where resource molecules (e.g. polymerases or ribosomes) bound DNA or mRNA molecules in a reversible reaction. Protein expression proceeded from the resource-DNA/RNA complex (Fig. 4B). The rate of protein expression (v) in this model is well-approximated by the Michaelis-Menten equation

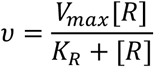

where *V*_*max*_is the maximum rate of protein production, [*R*] is the resource concentration, and *K*_*R*_ is the value of [R] where 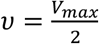. As [*R*] increases within a given expression center, the protein 3 production rate (i.e. the burst size) increases. However, there is diminishing marginal utility as each additional resource molecule added to the bursting center increases v by a smaller amount than the previously added resource molecule (Fig. 4C). The efficiency of a bursting center is

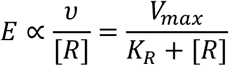

which monotonically decreases as [*R*] increases (Fig. 4D). So the net effect of a distribution of resources into a smaller number of bursting centers is to increase the burst size, but at the cost of decreasing expression efficiency. As a result, vesicles with only a few expression centers will result in low expression efficiency as it will force a high concentration of resources into each of the few centers. Conversely, vesicles with many expression centers will result in high expression efficiency as the resources are sparsely distributed across the many centers.

Experiments with macromolecular crowding agents report the direct observation of transcriptional expression centers and provide evidence that the centers nucleate and grow due to the hindered diffusion of mRNA and polysomes^9,10^. However, the total (i.e. including translation) expression behavior of these transcriptional centers is unknown, and it is unclear if, or how, such centers form in less crowded environments. However, in recent work we showed that increasing the population of expression resources leads to bigger burst sizes instead of the nucleation of new expression centers^8^. One possible interpretation of this result is that there is cooperativity at play in expression center formation, and that the capture of resources by a center makes the capture of additional resources more likely. Such positive feedback exists between transcription and translation with E. coli promoters^38^, and likely exists with T7 as well^9^.

The results presented here demonstrate that self-organization controls expression even more than the abundance of the molecular components of transcription and translation in confined cell-free gene expression. However, the corollary – that expression noise can be set by design – may be an equally important observation. In particular, expression noise is an essential feature in many probaBilistic fate-determining gene circuits^31,39,40^, and allows for quick adaptation to changing environments^39,41^. The independent control of expression mean and noise was accomplished through promoter design^42,43^ or with drugs^44^ in cell-Based systems. However, the central feature of cell-free synthetic biology is the ability to define the gene expression environment (e.g. confinement volume, degree of macromolecular crowding, composition of cell extract^45^, etc). Recent work^8–10^ demonstrated the use of confinement or macromolecular crowding to control expression noise. The results presented here demonstrate that manipulation of the concentration of expression resources may provide even finer control of expression noise.

Spatial organization is a defining attribute of cells and organisms, and may be as central to function as the genetic material^46^. Yet, cell-free synthetic biology has focused much more on specific molecular interactions that affect gene expression than on spatial considerations. Such an approach may have been appropriate for bulk reactions focused on maximizing protein production^47^. However, at cell-relevant levels of confinement, self-organization may lead to unanticipated and counterintuitive behavior. For example, here we showed that expression variation was largely unrelated to stochastic seeding of expression resources. Even more surprisingly, we found that the dilution of expression resources significantly reduced expression noise. Neither of these effects were primarily related to specific molecular interactions, but instead were sensitive to how spatial distribution of resources was changed by the concentration of resources. As cell-free systems are pushed to even higher levels of functional density, with the associated increase in macromolecular crowding and weak, non-specific interactions, more surprising behaviors are likely to emerge. Reaching the goal of cell-free synthetic biology that approaches cell-like capabilities will require uncovering and understanding these behaviors, and ultimately, to harness these behaviors to enable function.

## Methods

### Vesicle Preparation

Vesicle were made using the oil-in-water emulsion technique^48,49^ (Fig. 2A). This method encapsulated a protein expressing inner solution in vesicles separated from an osmotically balanced outer solution. The inner solution was prepared using 10 µL Solution A and 7.5 µL Solution B of the PURE system; 5 µL of sucrose solution (1 M); 0.25 µL of Transferrin-AlexaFluor 647; 0.125 µL of RNAsin (40 U/µL); 0.418 µL (1.67 nM) of YFP encoding pEToppYB plasmid^14^ (200 ng; 478.2 ng/µL); and nuclease-free water to bring the total volume of solution to 30 µL. The inner solution was vortexed in 330 µL of paraffin oil containing 30 mg of POPC for 60 seconds. The resulting emulsion was layered above the outer solution and centrifuged at 13,000 g for 20 minutes at room temperature.

### Outer Solution Preparation

The outer solution for vesicles was mixed from frozen stocks before each experiment. 1.5 µL Amino acid solution, 11.3 µL of ATP (100 mM), 7.5 µL of GTP (100 mM), 0.75 µL of CTP (500 mM), 0.75 µL of UTP (500 mM), 1.8 µL of spermidine (250 mM), 3.75 µL of creatine phosphate (1 M), 4.5 µL of Dithiothreitol (100 mM), 0.75 µL of Folinic Acid (0.5 M), 24 µL of potassium glutamate (3.5 M), 11.3 µL of magnesium acetate (0.5 M), 30 µL of HEPES (1 M), 60 µL of glucose (1 M), and 141.8 µL of autoclaved type I pure water for a total volume of 300 µL.

### Vesicle Imaging

The pellet of vesicles was collected with 100 µL of the outer solution and pipetted onto a no. 1.5 glass bottom petri dish. The lid was placed on the petri dish to minimize airflow and evaporation of the 100 µL outer solution and vesicle drop. The petri dish was placed on Zeiss LSM710 confocal scanning microscope with an incubation chamber warmed to 37° C and imaged every 3 minutes in a z-stack with a 20x air objective. YFP was excited with a 488 nm laser and fluorescent emission was collected from 515-584 nm. Similarly, AF647 was excited with a 633 nm laser and fluorescent emission was collected from 638-756 nm. Z-stacks were made of ∼21 slices at 1 µm intervals, and the aperture for each slice was 1.00 Airy Units (open enough to allow ∼1.5 µm depth of light). The time the vesicles sat on the microscope before imaging was minimized (less than 15 minutes), allowing for imaging for most of the duration of protein expression.

### Data Acquisition and Analysis

Average fluorescent intensity and diameter were measured with the FIJI TrackMate^50^ (v3.8.0) plugin. TrackMate found spots with an estimated bloB diameter of 10 µm using the Laplacian of Gaussian detector. Spots that were found with an estimated diameter <5 µm, >19 µm, or contrast <0 were removed from the data set. We used the simple Linear Assignment Problem (LAP) tracker to link spots across z-stacks in time to create traces. Traces that had missing frames, traveled >5µm between frames, or tracked for <45 of the 60 frames were removed from the data set. Fluorescent concentrations and estimated diameters were measured for each vesicle at each time point. The estimated diameter varied slightly between frames, so the diameter used for each vesicle was taken to be the average of the estimated diameters over the entire time trace. Protein abundance (population in arbitrary units) was found by multiplying the average fluorescent intensity by the volume. Temporal analysis of noise was done as described previously^8^. In short, fluorescent intensity time traces were corrected for background fluorescence. The deterministic trend of protein expression was removed from each trace to isolate the noise component. The coefficient of variation squared (CV^2^) was measured for each noise trace. We used the fluorescence value at the end of the experiment with equation 1 to calculate the burst size and frequency. We measured fluorescent signal from AF647 at the beginning of the experiment as a proxy for resource encapsulation statistics.

## Acknowledgements

This research was conducted at the Center for Nanophase Materials Sciences, which is a DOE Office of Science User Facility. P.M.C. and S.E.N. also acknowledge Graduate Fellowships from the bredesen Center for Interdisciplinary Research and Graduate Education, University of Tennessee, Knoxville. The authors would like to thank Osaka University and Dr. Tetsuya Yomo for providing pEToppYB plasmid.

